# Structure-guided, physics-aware reconstruction expands scan-limited multiphoton imaging

**DOI:** 10.64898/2026.04.20.719672

**Authors:** Shoupei Liu, Yaoguang Zhao, Jiahao Hu, Yanfeng Zhu, Huiyun Yu, Bo Li

**Affiliations:** School of Psychology, Institute of Chinese Cultural Psychology, Shanghai Jiao Tong University, Shanghai 200030, China; Huashan Hospital, Jinshan Hospital, MOE Frontiers Center for Brain Science, State Key Laboratory of Brain Function and Disorders, Institute for Translational Brain Research, Fudan University, Shanghai 200032, China; Wuhan Jianmin DAPENG Pharmaceutical Co., Ltd., Wuhan 430041, China; Department of Pharmacology, School of Basic Medical Sciences, Fudan University, Shanghai 200032, China

## Abstract

Multiphoton microscopy is increasingly pushed toward deep, large-scale and fast functional imaging, where acquisition speed has become a bottleneck. Here we present SPARC, a structure-guided, physics-aware reconstruction framework for scan-limited multiphoton imaging. Rather than treating undersampled recovery as generic super-resolution, SPARC formulates it as a constrained reconstruction problem tailored to point-scanning microscopy by incorporating a sample-matched structural reference from the same field of view and the anisotropic acquisition physics of scan-limited imaging. SPARC jointly integrates denoising and deep upsampling, enabling stable recovery from noisy sparse measurements. In simulated calcium imaging with ground truth, SPARC improved spatial reconstruction fidelity and temporal signal recovery relative to existing methods. *In vivo*, SPARC improved temporal readout in three-photon and mesoscale two-photon calcium imaging, and enabled a more temporally informative 400-Hz voltage readout on a standard resonant-galvo two-photon microscope. These results suggest SPARC expands the functional operating range of existing multiphoton microscopes under scan-limited conditions.

## Introduction

Multiphoton microscopy has transformed *in vivo* neuroscience by enabling optical access to neural activity deep inside intact brain tissue with cellular resolution^1–4^. As neuroscience moves toward deeper structures, larger-scale circuits and faster neural dynamics, multiphoton imaging is increasingly pushed beyond the operating regimes in which conventional high-resolution acquisition remains sufficient. Three-photon microscopy has opened functional imaging in deep cortical and subcortical structures^5–7^, mesoscale two-photon systems have extended cellular-resolution imaging across multiple brain regions^8–11^, and high-speed two-photon approaches are pushing optical recording toward direct measurements of membrane voltage^12–16^. Yet across these frontier applications, acquisition speed has emerged as a shared limiting factor.

This bottleneck arises for distinct physical reasons in each regime. In three-photon microscopy, the low repetition rates of excitation sources limit the effective pixel rate, such that functional imaging at 512 × 512 pixels is often restricted to frame rates of only ∼3–4 Hz^2,17–19^, which is insufficient for many fast indicators such as GCaMP6f. In mesoscale two-photon imaging, fields of view approaching ∼6 × 6 mm² require millions of sampling points per frame^11,20–22^, often reaching 4096 × 4096 pixels at cellular resolution, again driving frame rates below those needed for fast functional signals. For voltage imaging, the temporal requirement is even more stringent: resolving membrane potential dynamics can require ∼400 Hz sampling^23–25^, whereas a conventional 8-kHz resonant-scanning two-photon microscope reaches only ∼100 Hz even for a relatively small 128 × 512 field.

Microscopy reconstruction methods have advanced rapidly in recent years, progressing from supervised low-resolution-to-high-resolution inference^26–29^ to frameworks that incorporate temporal redundancy^27,28,30^, self-supervision^30–33^, and generic optical priors^31,32,34^. These advances show that computational reconstruction can extend the usable operating range of optical microscopes. Nevertheless, many existing formulations still face two general limitations. First, they recover high-resolution detail primarily by inferring it from undersampled measurements, making reconstruction fundamentally ill-posed and therefore prone to ambiguity and reconstruction artifacts^30–32,34^. Second, more aggressive undersampling leaves more information unmeasured, increasing the risk of hallucination and reconstruction instability^30–32,34^. These limitations suggest that, for scan-limited multiphoton imaging, reconstruction should not be formulated as generic detail inference from undersampled inputs alone.

For scan-limited multiphoton functional imaging, however, two constraints are often available that are absent from generic image reconstruction settings. First, a high-resolution structural image can often be acquired from the same sample and field of view before fast functional imaging, providing a specimen-matched spatial scaffold for recovery. Second, in many point-scanning multiphoton microscopes, acquisition time is limited primarily by sampling along the slow axis^22^, making anisotropic sparsification more physically realistic than isotropic downsampling under a matched imaging-time budget. Together, these considerations argue that scan-limited multiphoton reconstruction should be formulated as a structure-constrained, physics-aware inverse problem rather than generic super-resolution.

Here we introduce SPARC, a structure-guided, physics-aware reconstruction framework for scan-limited multiphoton imaging. Instead of treating undersampling as generic isotropic resolution loss, SPARC explicitly models the anisotropic acquisition constraints of point-scanning microscopy, in which imaging speed is often limited primarily by sampling along the slow axis. It further incorporates a sample-matched structural reference, acquired from the same field of view before fast imaging, to constrain recovery with specimen-specific spatial information rather than unconstrained inference alone. We validate this formulation in simulated calcium imaging with ground truth and then test it in three experimentally demanding regimes: deep three-photon calcium imaging, mesoscale two-photon calcium imaging, and high-speed two-photon voltage imaging. Across these settings, SPARC extends the effective spatiotemporal operating range of existing multiphoton microscopes without requiring new imaging hardware.

## Results

### SPARC establishes a structure-guided, physics-aware reconstruction framework for scan-limited multiphoton imaging

Existing microscopy reconstruction methods typically recover missing information by learning generic mappings from undersampled measurements, which makes the problem inherently ill-posed and susceptible to ambiguity and hallucination. For scan-limited multiphoton imaging, however, two additional constraints are often available but underused: a structural reference from the same specimen and field of view, and the anisotropic acquisition physics of point-scanning microscopes. SPARC is designed to exploit both.

For scan-limited multiphoton imaging, a particularly informative constraint is often available before fast functional acquisition begins. Unlike generic image reconstruction tasks, multiphoton experiments often allow a low-frame-rate but high-spatial-resolution full-frame reference image to be acquired from the same sample and field of view (Fig. 1a). Although this structural reference is not a frame-by-frame ground truth for the subsequent dynamic sequence, it provides a specimen-matched spatial scaffold that constrains recovery with actual specimen structure rather than unconstrained inference of missing details. We therefore formulated SPARC as a structure-guided recovery framework that explicitly incorporates this sample-specific structural information into reconstruction. Importantly, the structural reference does not provide frame-specific functional information; instead, it constrains spatial recovery with specimen-matched anatomy, while temporal fluctuations remain determined by the sparse functional measurements.

**Fig. 1.**
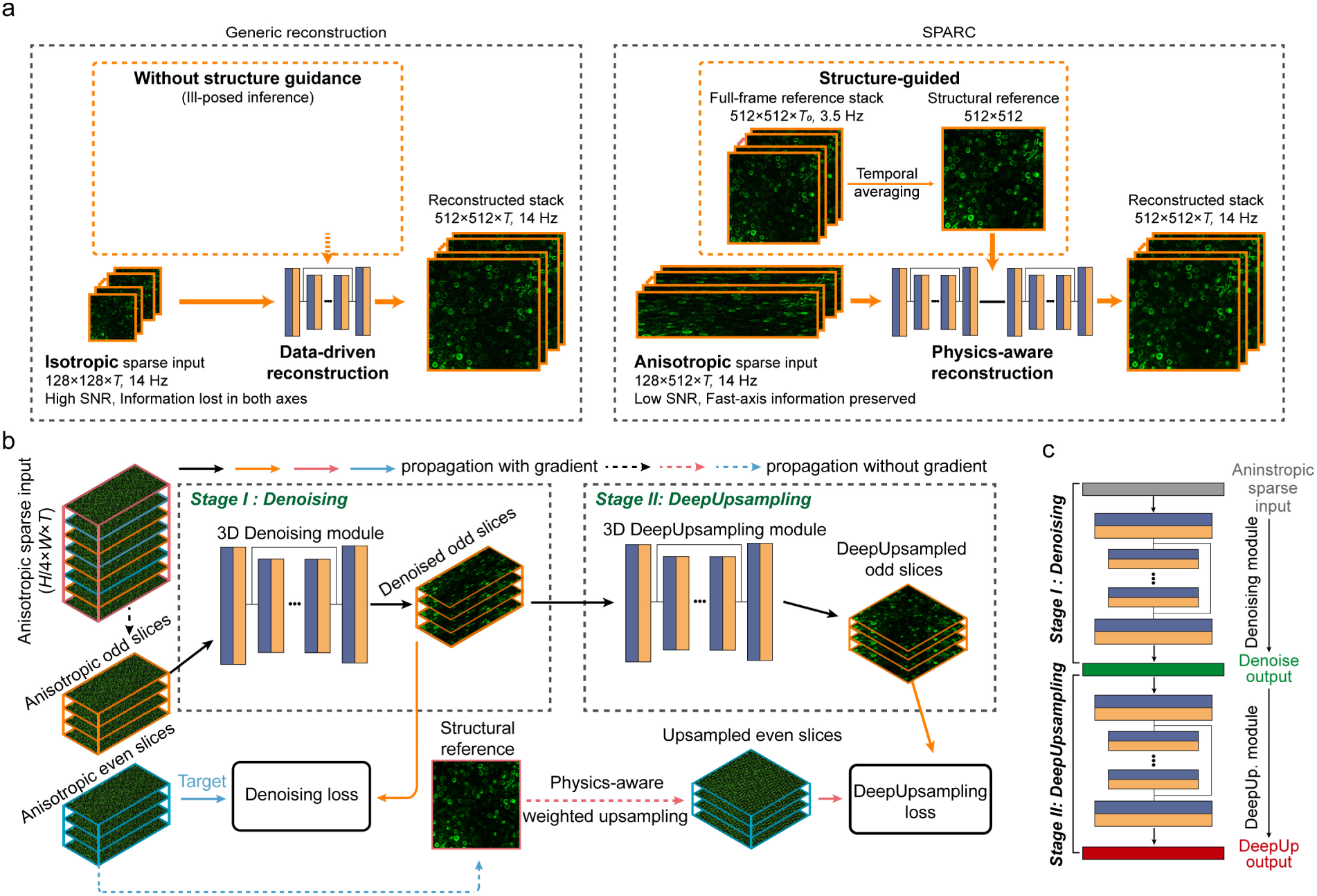
SPARC framework integrates structure-guided, physics-aware reconstruction with anisotropic sampling. **a**, Conceptual comparison between generic reconstruction and SPARC. Left, generic reconstruction takes an isotropic sparse input and performs data-driven reconstruction of a high-resolution image stack without structural guidance. Right, SPARC takes an anisotropic sparse input and reconstructs the high-resolution stack using sample-specific structural guidance and physics-aware reconstruction. A structural reference is obtained by temporally averaging a low-frame-rate full-frame reference stack. **b**, Training workflow of the SPARC two-stage network. Stage I performs denoising of anisotropic sparse input frames, producing a denoised sparse stack. Stage II then performs upsampling guided by the structural reference and physics-aware weighted upsampling. The two stages are jointly optimized using denoising loss and deep-upsampling loss. **c**, Inference workflow of SPARC. The sparse anisotropic input is first processed by the denoising module and then by the deep-upsampling module to generate the reconstructed output.

We next asked how undersampling should be defined under the scan physics of point-scanning multiphoton microscopy (Fig. 1a). In many practical systems, total acquisition time is determined mainly by the number of scan lines along the slow axis, whereas sampling density along the fast axis can often be preserved without changing the overall imaging-time budget. Under a matched imaging-time budget, sparsifying only the slow axis while preserving full fast-axis sampling is therefore more physically realistic than isotropic downsampling. This anisotropic strategy retains more directly measured fast-axis information and thus preserves more recoverable information for subsequent reconstruction. However, this advantage does not automatically translate into better recovery. Under a fixed line period, preserving full fast-axis sampling distributes the available signal over more fast-axis pixels and can reduce the effective input signal-to-noise ratio (SNR). As a result, anisotropic inputs outperform isotropic ones under ideal low-noise conditions (Supplementary Fig. 1), but in typical calcium imaging settings, where fast functional movies are noisier, the lower input SNR can outweigh the benefit of preserving more measured information. Thus, anisotropic sampling preserves more recoverable information, but by itself does not guarantee better reconstruction under noisy conditions.

This trade-off motivates denoising before upsampling. However, when a reconstruction task requires both denoising and upsampling, treating them as two separately optimized procedures can be suboptimal^34^. A denoising module trained in isolation typically aims to produce a generally cleaner image, rather than a representation specifically optimized for subsequent structure-guided, physics-aware upsampling. As a result, weak structural signals that remain useful for reconstruction may be suppressed, and biases introduced during denoising may become fixed and propagate into the following upsampling stage. To address this issue, we designed SPARC as a unified two-stage deep network in which denoising and upsampling are integrated rather than independently optimized (Fig. 1b, Supplementary Fig. 2). In Stage I, the anisotropic sparse input is denoised to stabilize the recoverable measurement content. In Stage II, the denoised sparse stack is upsampled under the joint guidance of the sample-specific structural reference and physics-aware weighted upsampling (Supplementary Fig. 3) to restore the missing slow-axis information. The network parameters are optimized jointly through backpropagation from the denoising and deep-upsampling losses, allowing the first stage to preserve information in a way that directly serves the final recovery objective. During inference, the sparse anisotropic input is first processed by the denoising module and then by the deep-upsampling module to generate the reconstructed output (Fig. 1c). In this way, SPARC unifies noise suppression, anisotropic sampling, and structure-guided, physics-aware recovery in a single framework for scan-limited multiphoton imaging. Conceptually, SPARC is therefore not a generic super-resolution network with an added reference input, but a reconstruction framework tailored to the acquisition logic of point-scanning multiphoton imaging.

### SPARC improves spatial and temporal fidelity in ground-truth calcium imaging benchmarks

To benchmark SPARC in a controlled setting with known ground truth, we generated realistic two-photon calcium imaging data using the Neural Anatomy and Optical Microscopy (NAOMi)^35^ simulator. To emulate practical calcium imaging conditions, mixed Gaussian and Poisson noise were added to generate a raw noisy image stack with an SNR of 5 dB (512 × 512 × 1008 frames, 14 Hz). Under matched acquisition-time constraints, sparse input stacks were generated using isotropic and anisotropic downsampling (isotropic, 128 × 128 × 1008; anisotropic, 128 × 512 × 1008), while maintaining the same temporal sampling rate. Because conventional reconstruction did not benefit from anisotropic input under this noisy calcium-imaging regime (Supplementary Fig. 1), we used the conventional isotropic setting for the existing baseline methods (Self-vision^33^ and DFCAN^29^) and focused the benchmark on whether SPARC could specifically exploit anisotropic sparse acquisition under scan-limited conditions (Fig. 2a).

**Fig. 2.**
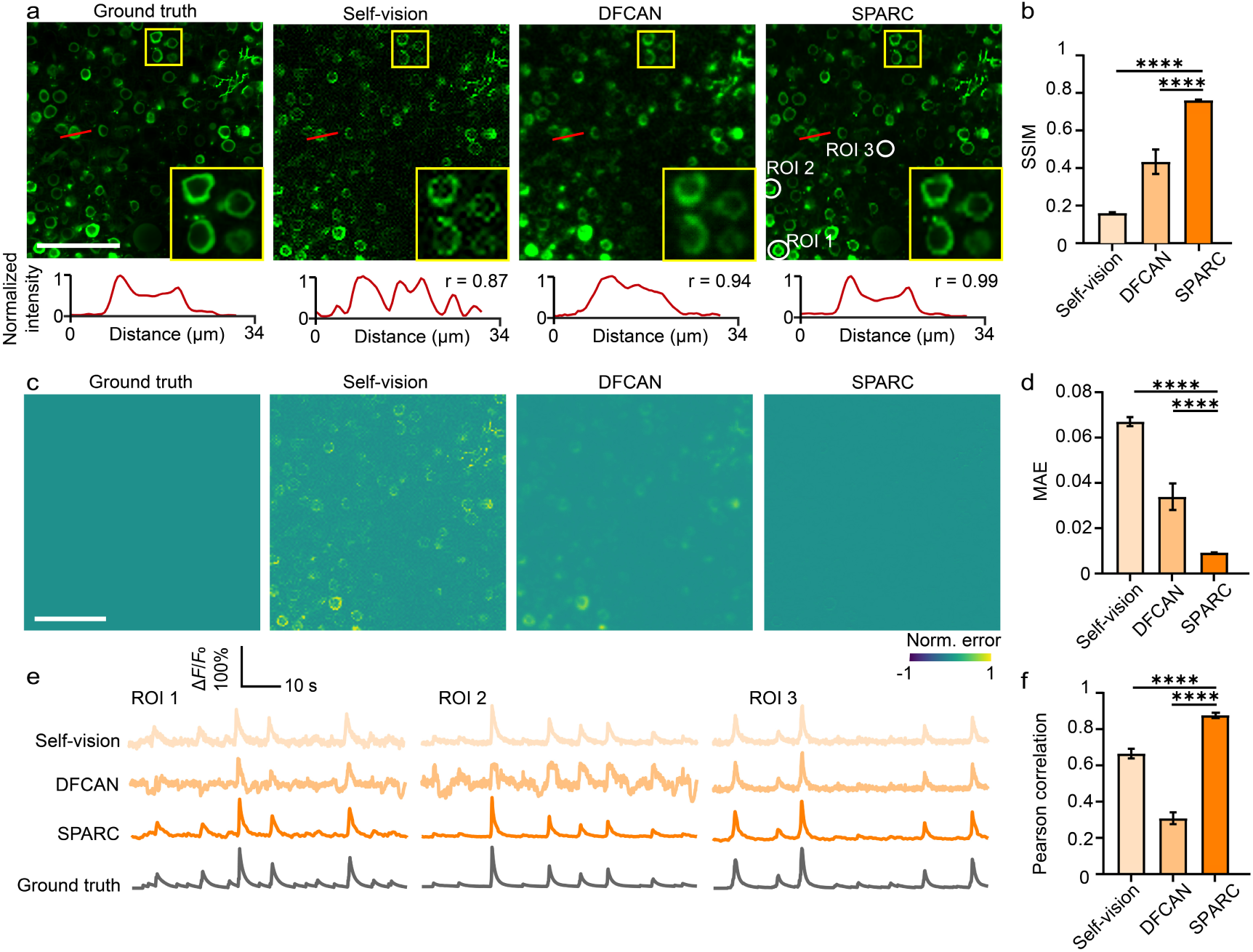
Performance validation of SPARC on simulated calcium imaging data. **a**, Representative single-frame images reconstructed from calcium imaging data sampled at 14 Hz. From left to right: ground truth, Self-vision, DFCAN, and SPARC. Yellow boxes show magnified views of the indicated regions. The bottom row shows normalized intensity profiles along the red lines, with Pearson correlation coefficients (r) relative to the ground truth indicated for each reconstruction. Scale bar, 100 μm. **b**, Structural similarity index (SSIM) of Self-vision, DFCAN, and SPARC relative to the ground truth. Error bars indicate s.d. Statistical significance was assessed using paired *t*-test with *n* = 1008 image frames per group. \*\*\*\**P* < 0.0001. **c**, Normalized error maps of ground truth, Self-vision, DFCAN, and SPARC relative to the ground truth. Pixel values are displayed over the range [-1, 1]. Scale bar, 100 μm. **d**, Mean absolute error (MAE) of Self-vision, DFCAN, and SPARC relative to the ground truth. Error bars indicate s.d. Statistical significance was assessed using paired *t*-test with *n* = 1008 image frames per group. \*\*\*\**P* < 0.0001. **e**, Calcium activity traces from three regions of interest (ROIs) indicated by the white circles in **a**. Rows correspond to Self-vision, DFCAN, SPARC, and ground truth. **f**, Pearson correlation coefficients of neuronal calcium activity traces relative to the ground truth. Error bars indicate s.e.m. Statistical significance was assessed using paired *t*-test with *n* = 92 neurons. \*\*\*\**P* < 0.0001.

Qualitatively, SPARC reconstructed neuronal morphology more clearly than the comparison methods, preserving finer processes and local microstructures that were less distinct in the other reconstructions (Fig. 2a). This improvement was also evident in the intensity profiles along the selected line cuts, where the SPARC reconstruction most closely matched the ground truth, with the highest profile correlation (r = 0.99; Fig. 2a). We next quantified frame-wise spatial fidelity using the structural similarity index (SSIM). SPARC achieved the highest SSIM among all methods (0.7609 ± 0.0039, mean ± s.d.), indicating more accurate recovery of spatial structure relative to the comparison methods (Fig. 2b). Consistent with this result, normalized error maps showed that SPARC had the smallest residual errors across the field of view, indicating more accurate recovery of the underlying calcium imaging data (Fig. 2c). This result was further supported by the mean absolute error (MAE), which was the lowest for SPARC among all methods (0.0092 ± 0.0001, mean ± s.d.; Fig. 2d).

We next asked whether the improved image reconstruction also translated into more faithful recovery of neuronal activity over time. Calcium traces extracted from representative neurons showed that SPARC more closely followed the ground-truth dynamics while exhibiting less fluctuation noise than the other methods (Fig. 2e). Across 92 neurons, Pearson correlation analysis demonstrated that SPARC achieved the highest trace-level agreement with the ground truth (0.8767 ± 0.0125, mean ± s.e.m.), significantly exceeding the comparison methods (Fig. 2f). Together, these results indicate that SPARC improves both spatial reconstruction quality and temporal signal fidelity in simulated calcium imaging benchmarks with synchronized ground truth.

### SPARC increases the effective temporal resolution of *in vivo* three-photon calcium imaging

Three-photon microscopy has greatly extended the accessible depth of *in vivo* calcium imaging, enabling functional recordings from deep cortical and subcortical circuits that are difficult to reach with conventional two-photon^5,6,36^ excitation. This capability is particularly important for studying neural dynamics in physiologically intact deep-brain networks^2,17,37–39^. However, the temporal resolution of three-photon calcium imaging remains fundamentally constrained by the imaging physics. Because three-photon excitation requires high pulse energies, most practical systems operate at relatively low laser repetition rates, often around 1 MHz. Under these conditions, the achievable pixel rate is intrinsically limited, such that conventional full-frame imaging at 512 × 512 pixels is typically restricted to only a few hertz (< 4 Hz^2,17–19^). While such frame rates may be adequate for relatively slow calcium indicators, they increasingly limit the ability to resolve faster calcium dynamics as indicator kinetics continue to improve. We therefore asked whether SPARC could alleviate this temporal bottleneck by reconstructing high-resolution, high-frame-rate image sequences from anisotropically sampled three-photon data under the same fast functional acquisition budget.

To test this idea, we performed *in vivo* three-photon calcium imaging in layer VI of the mouse sensory cortex expressing GCaMP6f. The same field of view was acquired using two complementary imaging modes. First, a conventional full-frame acquisition mode (512 × 512, 3.38 Hz) was used to obtain a low-frame-rate, high-resolution image stack, from which 1,000 frames were averaged to generate the structural reference image (Fig. 3a, left). Second, a high-speed anisotropic sampling mode (128 × 512, 13.52 Hz) was used to acquire sparse measurements at a fourfold higher frame rate (Fig. 3a, middle). These anisotropically sampled data were then reconstructed by SPARC, yielding high-resolution image sequences at the same high temporal sampling rate (512 × 512, 13.52 Hz; Fig. 3a, right).

**Fig. 3.**
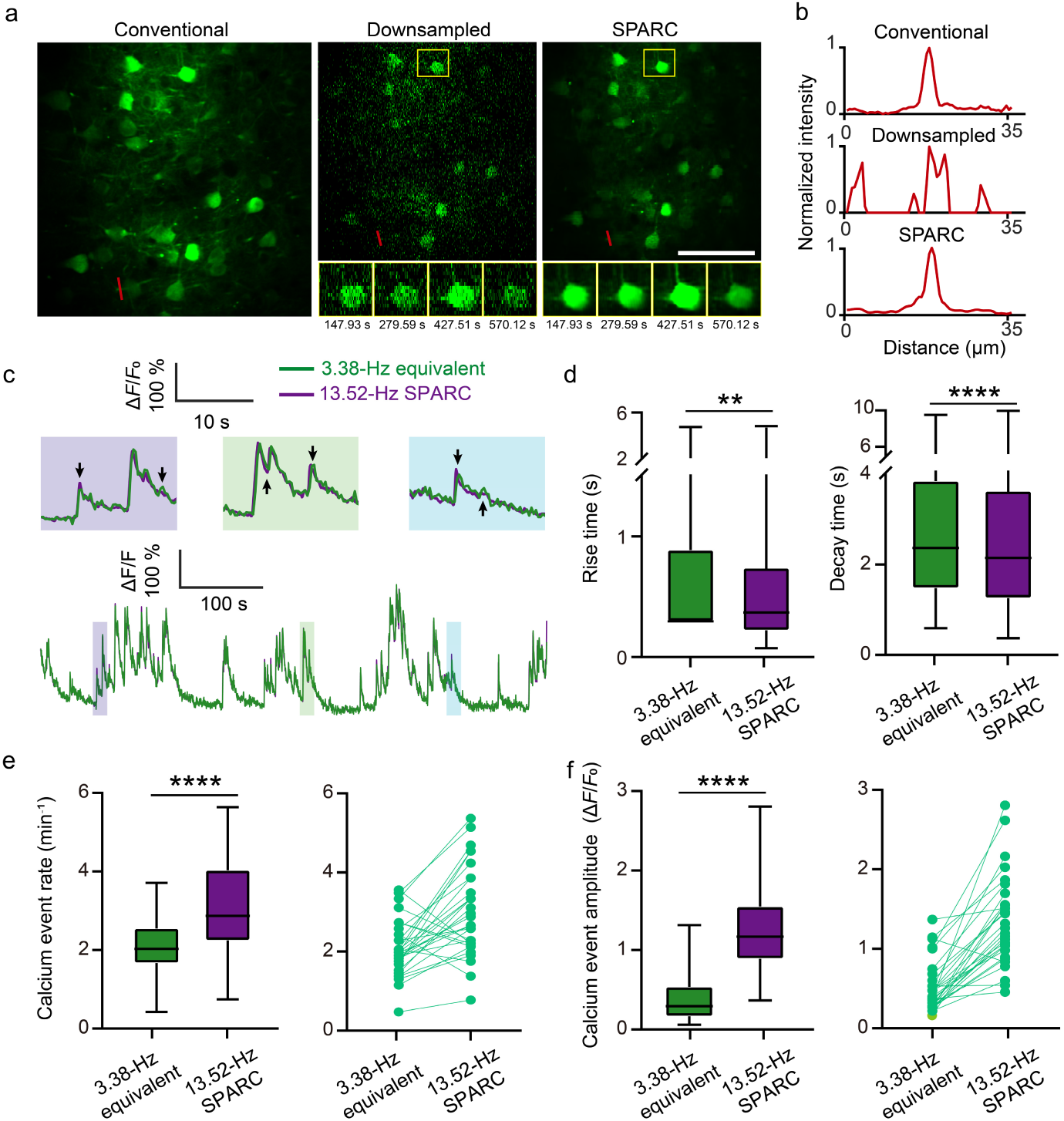
SPARC improves the effective temporal resolution of *in vivo* three-photon calcium imaging. **a**, Representative three-photon calcium imaging results from the same field of view in a GCaMP6f-labeled mouse. Left, conventional full-frame image (512 × 512, 3.38 Hz, 1,000 frames averaged). Middle, anisotropically downsampled single frame (128 × 512, 13.52 Hz). Right, SPARC single frame (512 × 512, 13.52 Hz). Scale bar, 100 μm. **b**, Normalized intensity profiles along the red lines in **a**. **c**, Representative calcium activity traces comparing the 3.38-Hz equivalent (green; obtained by temporal downsampling of the anisotropic sparse stack) and the 13.52-Hz SPARC signal (purple). Insets show magnified views of calcium transients indicated by the shaded boxes. **d**, The apparent rise time (left) and decay half-time (right) of calcium transients for the 3.38-Hz equivalent and the 13.52-Hz SPARC signal (*n* = 586 matched events from 30 neurons). **e**, Left, calcium event rate for the 3.38-Hz equivalent and the 13.52-Hz SPARC signal (*n* = 30 neurons). Right, paired changes in event rate for individual neurons. **f**, Left, calcium event amplitude for the 3.38-Hz equivalent and the 13.52-Hz SPARC signal (*n* = 30 neurons). Right, paired changes in event amplitude for individual neurons. Statistical significance was assessed using a paired *t*-test. \*\**P* = 0.0027, \*\*\*\**P* < 0.0001.

At the single-frame level, SPARC substantially improved image quality relative to the raw anisotropic input, recovering neuronal morphology and fine processes that were difficult to resolve in the sparse measurements alone (Fig. 3a). This improvement was consistently evident across representative time points, where SPARC restored structural details that were blurred or fragmented in the raw sparse frames. Line-profile analysis further showed that the reconstructed spatial intensity patterns more closely matched those of the structural reference image, indicating that SPARC preserved structural fidelity while increasing the effective frame rate (Fig. 3b).

We next asked whether this gain in spatiotemporal reconstruction translated into a more informative functional readout over time. Here, the key question was whether structure-guided reconstruction could convert this faster but sparse acquisition into a functionally more informative readout. To address this, we compared calcium traces from the 13.52-Hz SPARC reconstruction with a 3.38-Hz equivalent signal obtained by temporal downsampling of the sparse data. Over a 600-s recording period, the higher-frame-rate SPARC traces more clearly resolved calcium transient kinetics, including steeper rising phases, sharper peaks, and more distinct decay trajectories (Fig. 3c). Quantitative analysis further showed that SPARC improved multiple kinetic and event-level readouts relative to the equivalent low-frame-rate signal. In matched-event comparisons, the apparent rise time and decay half-time were both significantly shortened, indicating more precise temporal delineation of transient dynamics (Fig. 3d). In addition, SPARC increased the detected event rate and yielded larger measured event amplitudes (Fig. 3e,f), consistent with improved detectability of fast and transient calcium activity. Together, these results indicate that SPARC can provide a more temporally informative readout of *in vivo* three-photon calcium imaging while preserving high spatial resolution under the tested acquisition conditions.

### SPARC increases the effective temporal resolution of mesoscale *in vivo* two-photon calcium imaging

Understanding brain function increasingly requires monitoring neuronal activity across distributed cortical networks rather than within a single local microcircuit. Mesoscale two-photon imaging has therefore emerged as an important approach for extending cellular-resolution calcium imaging across multiple millimeters of cortex. However, this regime introduces a distinct speed bottleneck^9,40^. When the field of view expands while single-cell resolution is maintained, the number of sampling points required per frame increases dramatically, often reaching millions of pixels. In practice, fields of view approaching 6 × 6 mm² can require sampling densities on the order of 4096 × 4096 pixels at cellular resolution, which substantially reduces the achievable frame rate under sequential laser scanning. We therefore asked whether SPARC could alleviate this mesoscale imaging bottleneck by preserving structural detail while increasing the effective temporal resolution of large-field-of-view two-photon calcium imaging. Here again, the key question was whether SPARC could improve temporal readout under the same fast functional acquisition budget. This setting provides a conceptually distinct test case from three-photon imaging, because here the limiting factor is not deep-imaging pixel rate but the massive sampling burden imposed by a large cellular-resolution field of view.

To first assess whether SPARC remained effective for mesoscale structural imaging, we performed two-photon imaging over a 6 × 6 mm² cortical region in a transgenic mouse expressing GCaMP6s in cortical excitatory neurons. A conventional full-frame acquisition mode (4096 × 4096, 3.8 Hz) was used to obtain a high-resolution image stack, from which 1,000 frames were averaged to generate the structural reference image. The same field of view was then imaged using a high-speed anisotropic sampling mode (1024 × 4096, 15.21 Hz) to acquire sparse measurements at a fourfold higher frame rate. SPARC reconstructed these sparse measurements into high-resolution image sequences at the same high frame rate (4096 × 4096, 15.21 Hz; Fig. 4a). At the single-frame level, SPARC restored fine neuronal morphology and vascular detail that were not clearly resolved in the anisotropically sampled input. This improvement was evident in magnified views of selected regions and was further supported by line-profile analysis, in which the reconstructed spatial intensity distributions more closely matched those of the structural reference image (Fig. 4b,c).

**Fig. 4.**
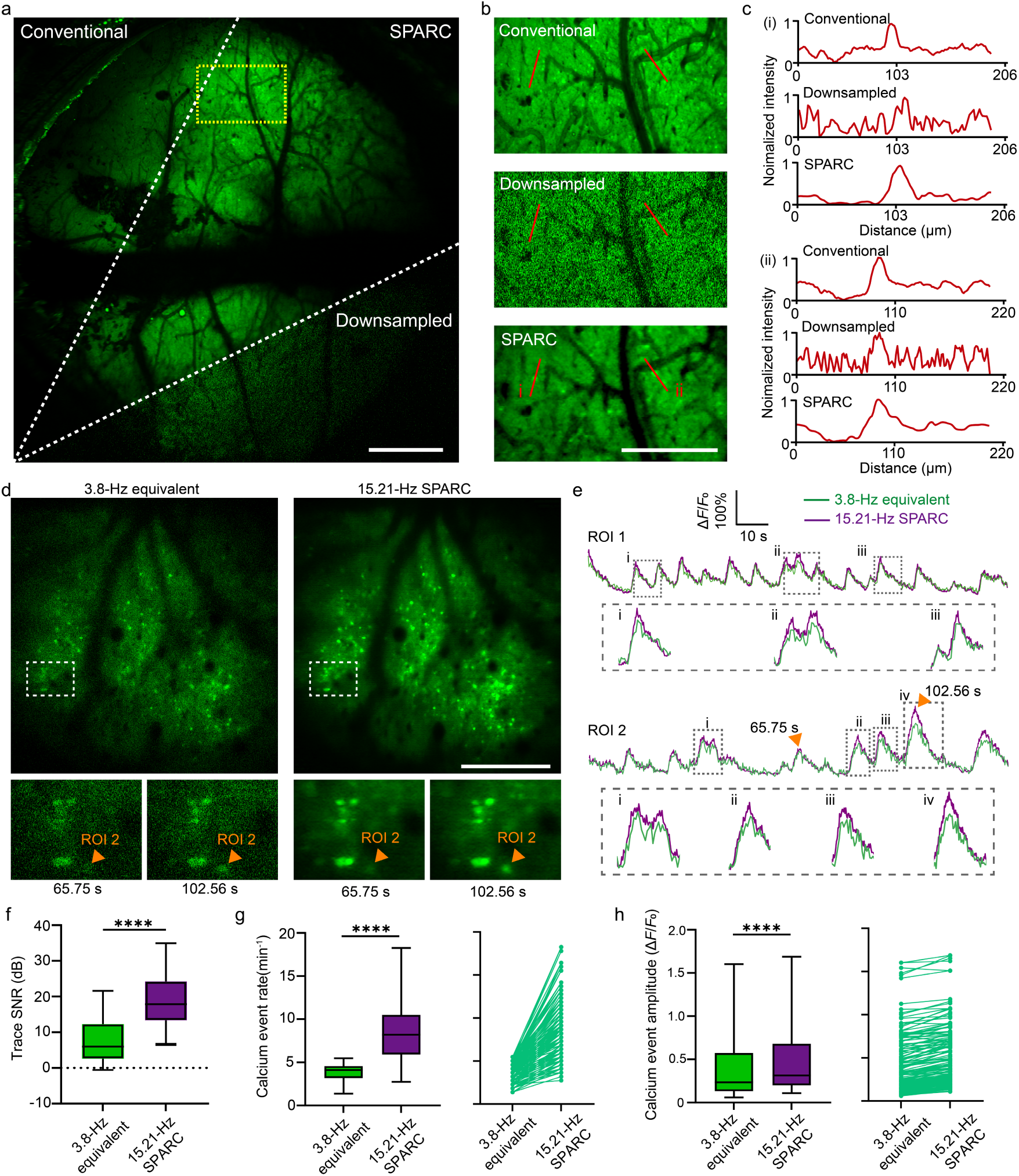
SPARC improves the effective temporal resolution of mesoscale *in vivo* two-photon calcium imaging. **a**, Representative mesoscale two-photon images (6 × 6 mm^2^) from a CaMKIIα::Ai162 transgenic mouse. Left, conventional full-frame image (4096 × 4096, 3.8 Hz, 1,000 frames averaged). Middle, SPARC single frame (4096 × 4096, 15.21 Hz). Right, anisotropically downsampled single frame (1024 × 4096, 15.21 Hz). Scale bar, 1 mm. **b**, Magnified views of the boxed region in **a**. Scale bar, 500 μm. **c**, Normalized intensity profiles along the two red lines in **b**. For each profile, traces are shown from top to bottom for the conventional full-frame image, downsampled input, and SPARC image. **d**, Representative high-speed mesoscale two-photon calcium imaging (1.5 × 1.5 mm^2^, 15.21 Hz) of GCaMP6f-labeled neurons from the same field of view in a virus-injected mouse. Left, 3.8-Hz-equivalent image obtained by temporal downsampling of the anisotropic sparse stack. Right, 15.21-Hz SPARC image. Bottom, magnified views of the boxed region at 65.75 s and 102.56 s. Scale bar, 500 μm. **e**, Representative calcium activity traces from two ROIs comparing the 3.8-Hz equivalent (green) and the 15.21-Hz SPARC signal (purple). Insets show magnified views of calcium transients indicated by dashed boxes. In ROI 2, the triangle marks the neuron whose fluorescence changes at 65.75 s and 102.56 s are highlighted in **d**. **f**, Trace signal-to-noise ratio (SNR) for the 3.8-Hz equivalent and the 15.21-Hz SPARC signal. The dashed line indicates SNR = 0 dB. **g**, Left, calcium event rate for the 3.8-Hz equivalent and the 15.21-Hz SPARC signal. Right, paired changes in event rate for individual neurons. **h**, Left, calcium event amplitude for the 3.8-Hz equivalent and the 15.21-Hz SPARC signal. Right, paired changes in event amplitude for individual neurons. Statistical significance was assessed using paired *t*-test (*n* = 182 neurons). \*\*\*\**P* < 0.0001.

We next asked whether this gain in spatiotemporal reconstruction translated into a more informative functional readout over time. To test this, we performed functional imaging over a 1.5 × 1.5 mm² field of view in mice expressing the faster calcium indicator GCaMP6f. We compared the 15.21-Hz SPARC reconstruction with a 3.8-Hz equivalent signal obtained by temporal downsampling of the anisotropic sparse stack (Fig. 4d). In representative frames, the SPARC reconstruction provided clearer single-frame visualization of neuronal calcium activity at matched time points. Consistently, calcium traces extracted from representative ROIs showed better-defined rise phases, peaks, and decay trajectories in the 15.21-Hz SPARC signal than in the 3.8-

Hz equivalent signal (Fig. 4e). Quantitative analysis confirmed these improvements at the population level. Relative to the 3.8-Hz equivalent signal, SPARC significantly increased trace signal-to-noise ratio, event detection rate, and event amplitude across recorded neurons (Fig. 4f–h). Together, these results indicate that SPARC can provide a more temporally informative readout of mesoscale two-photon imaging under the tested acquisition conditions.

### SPARC enables a more temporally informative 400-Hz voltage readout on a standard resonant-galvo two-photon microscope

Voltage imaging provides a uniquely direct optical readout of neuronal membrane potential, enabling action potentials, subthreshold fluctuations, and fast population synchrony to be measured at cellular resolution with substantially higher temporal precision than calcium imaging^15,24,25,41,42^. This capability is particularly important for studying circuit dynamics that unfold on millisecond timescales^43–45^. Recent advances in genetically encoded voltage indicators, including ASAP5, have further increased the demand for high-speed optical readout by improving the detectability of both spiking and subthreshold signals. However, these fast signals require sampling in the hundreds-of-hertz to kilohertz range, where two-photon voltage imaging becomes severely constrained by point-scanning speed and photon budget. As a result, *in vivo* two-photon voltage imaging has often relied on specialized high-speed optical implementations rather than standard resonant-galvo microscopes^12,15,16,46,47^. We therefore asked whether SPARC could provide a more temporally informative 400-Hz *in vivo* voltage readout on a standard resonant-galvo two-photon microscope under scan-limited acquisition.

The experimental configuration is schematized in Fig. 5a. We first acquired a low-frame-rate, high-spatial-resolution structural image of ASAP5-labeled neurons in the mouse somatosensory cortex using a conventional two-photon configuration (120 × 512 pixels per frame, 100 Hz, averaged over 10,000 frames; Fig. 5b, top). We then performed high-speed voltage imaging in the same field using anisotropic sparse sampling at 400 Hz (30 × 512; Fig. 5b, middle). Under these acquisition conditions, the raw single-frame images were strongly degraded by low pixel resolution and noise, making neuronal structure difficult to interpret directly. SPARC reconstructed the same data into high-resolution voltage images at 400 Hz (120 × 512; Fig. 5b, bottom), recovering neuronal morphology with markedly improved clarity.

**Fig. 5.**
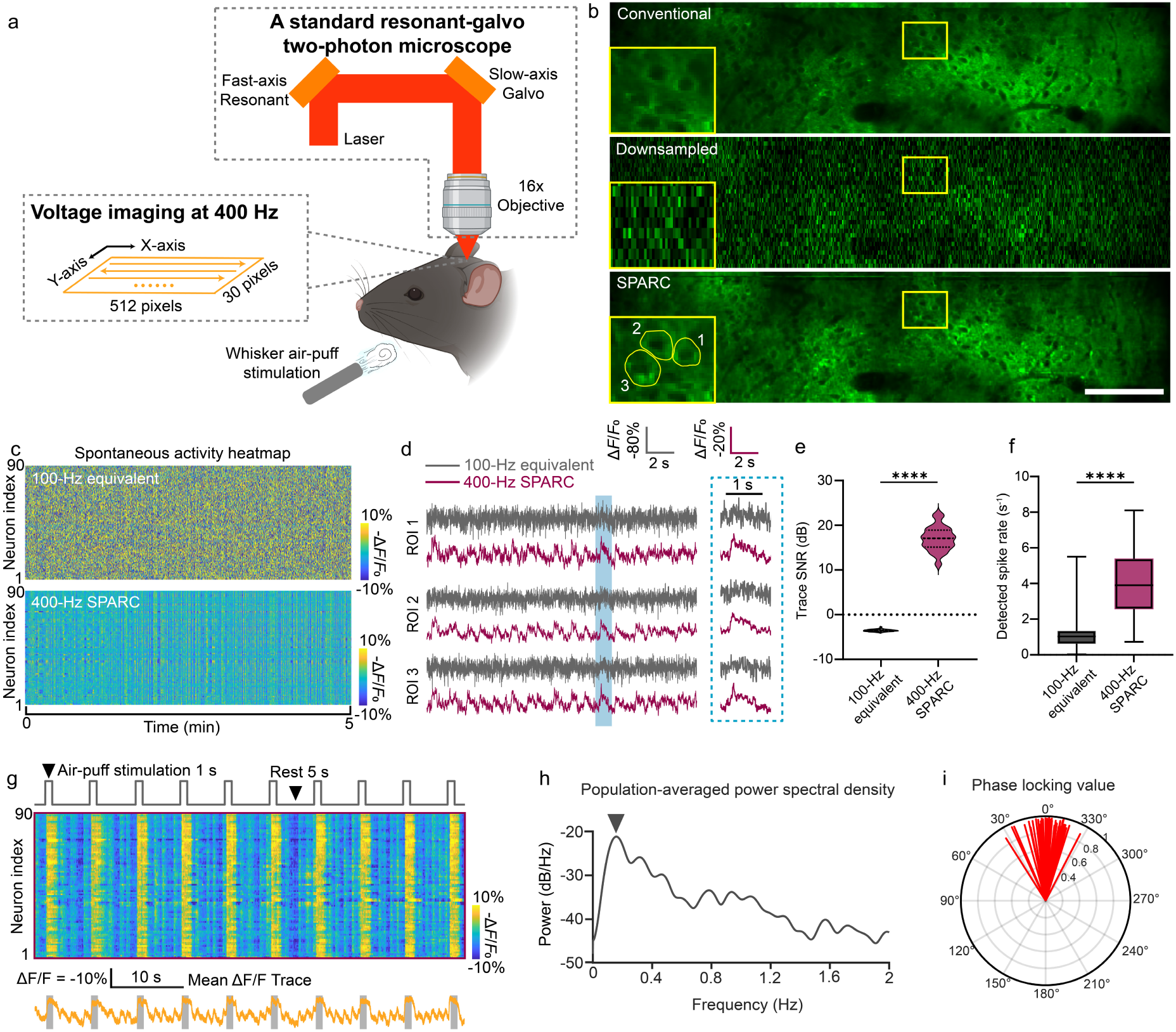
SPARC enables a more temporally informative 400-Hz voltage readout on a standard resonant-galvo two-photon microscope. **a**, Schematic illustration of the SPARC-supported two-photon imaging configuration and experimental paradigm for *in vivo* voltage imaging. With SPARC, a standard resonant-galvo two-photon microscope configuration can provide a more temporally informative 400-Hz voltage readout under scan-limited acquisition. **b**, Representative *in vivo* voltage imaging results from ASAP5-labeled neurons in the same mouse. From top to bottom: conventional full-frame image (120 × 512, 100 Hz, 10,000 frames averaged), anisotropically downsampled single frame (30 × 512, 400 Hz), and SPARC single frame (120 × 512, 400 Hz). Insets show magnified views of the boxed regions. Scale bar, 100 μm. **c**, Spontaneous activity heatmaps from 90 neurons comparing the 100-Hz equivalent (top; obtained by temporal downsampling of the anisotropic sparse stack) and the 400-Hz SPARC signal (bottom). **d**, Representative spontaneous voltage traces from three neurons comparing the 100-Hz equivalent (gray) and the 400-Hz SPARC signal (magenta). The dashed box at right shows magnified views of the traces within the shaded time window. The three neurons are indicated by the yellow outlines in **b**. **e**, Trace signal-to-noise ratio (SNR) of neuronal voltage traces for the 100-Hz equivalent and the 400-Hz SPARC signal. The dashed line indicates SNR = 0 dB. **f**, Detected spike rate for the 100-Hz equivalent and the 400-Hz SPARC signal. Statistical significance was assessed using paired *t*-test with *n* = 90 neurons. \*\*\*\**P* < 0.0001. **g**, SPARC reveals whisker air-puff-evoked synchronized population activity. Top, stimulation timeline in an awake head-fixed mouse. Middle, heatmap of population voltage activity from 90 neurons during periodic air-puff stimulation. Bottom, mean Δ*F*/*F*_0_ trace across the recorded population. **h**, Population-averaged power spectral density of the voltage activity shown in **g**. **i**, Polar distribution of phase-locking values for population voltage activity during periodic whisker air-puff stimulation. Voltage traces were filtered in the 0.1-10 Hz range.

We next evaluated whether SPARC could recover spontaneous voltage dynamics with improved temporal fidelity. We compared a 100-Hz equivalent signal, obtained by temporal downsampling of the anisotropic sparse stack, with the 400-Hz SPARC reconstruction within the recorded 90-neuron population. This comparison therefore asked whether structure-guided reconstruction could convert the same sparse acquisition into a temporally more informative readout rather than simply benefiting from longer acquisition or different hardware. The reconstructed activity heatmap more clearly resolved fluctuations in membrane potential and synchronous population activity (Fig. 5c). Representative traces from three neurons further showed that SPARC improved the recoverability of spike-like fast voltage transients and rendered their temporal structure more clearly detectable under the current analysis pipeline (Fig. 5d). Quantification further showed that the 400-Hz SPARC signal achieved significantly higher trace SNR and a markedly higher detected spike rate than the 100-Hz equivalent signal, with trace SNR increasing from −3.557 ± 0.0231 dB to 17 ± 0.2828 dB and detected spike rate increasing from 1.157 ± 0.0862 s⁻¹ to 4.065 ± 0.1995 s⁻¹ (mean ± s.e.m.; Fig. 5e,f).

Here we focused on population-level temporal organization as an additional validation dimension beyond single-trace visual quality. We further asked whether SPARC could resolve stimulus-coupled fast population dynamics. To test this, we applied periodic air-puff stimulation to the whiskers of awake head-fixed mice while recording voltage activity. Each cycle consisted of 1 s of air puff followed by 5 s of rest and was repeated over 10 cycles (Fig. 5g, top). The SPARC voltage activity heatmap revealed stimulus-locked synchronous population activity across the 90-neuron population (Fig. 5g, middle), and the population-averaged voltage trace closely followed the stimulation paradigm (Fig. 5g, bottom). Spectral analysis of the population-averaged trace revealed a clear low-frequency component aligned with the stimulation cycle (Fig. 5h). Phase-locking analysis further showed a tightly concentrated phase distribution within the 0.1–10 Hz band, indicating strong entrainment of population voltage activity to the sensory stimulus (Fig. 5i). Together, these results indicate that SPARC can provide a more temporally informative 400-Hz voltage readout on a standard resonant-galvo two-photon microscope under the present scan-limited acquisition conditions, while improving the recoverability of both spontaneous and stimulus-evoked fast population dynamics.

## Discussion

In this study, we developed SPARC, a structure-guided, physics-aware reconstruction framework for scan-limited multiphoton imaging. In simulated calcium imaging with ground truth, SPARC improved both spatial and temporal reconstruction fidelity relative to existing methods. Across three representative *in vivo* application settings, SPARC improved the effective spatiotemporal readout of scan-limited multiphoton imaging under the tested acquisition conditions. Together, these results suggest that a common acquisition-speed bottleneck shared by several major directions of multiphoton neural imaging may, in some cases, be alleviated computationally without requiring a distinct imaging platform for each scan-limited regime.

The performance of SPARC likely arises from the combination of three design features. First, the structural reference constrains recovery with sample-specific spatial correspondence rather than unconstrained detail inference. Second, anisotropic sampling preserves more directly measured information under a matched acquisition-time budget by retaining dense sampling along the fast axis while sparsifying the slow axis. Third, denoising and deep upsampling are optimized jointly rather than as isolated steps, allowing noise suppression to better serve the final reconstruction objective. Together, these features help distinguish SPARC from microscopy reconstruction approaches that rely primarily on data-driven inference from undersampled measurements alone.

SPARC also has important boundary conditions. First, it assumes that a structural reference can be acquired from the same field of view before fast functional imaging. This condition is often practical in multiphoton experiments, but it may be less suitable for preparations with rapid structural drift, substantial motion, or unstable long-term registration. Second, the current formulation is best matched to scan-limited point-scanning systems in which substantial speed gains can be achieved by sparsifying the slow axis while preserving dense sampling along the fast axis. Extension to substantially different acquisition geometries will likely require reformulating both the sampling model and the reconstruction target. Third, although the present results support improved recoverability of functional signals, reconstruction-based approaches should be interpreted with care, particularly for fast and low-SNR signals such as voltage imaging, where potential bias or hallucination must be carefully considered. The present *in vivo* analyses were designed to test algorithmic behavior across three representative scan-limited imaging regimes, rather than to estimate biological variability across animals. Accordingly, broader validation across additional animals, sessions, probes, preparations, imaging conditions and microscope architectures will be important for defining the full scope of generalizability of the framework.

Taken together, SPARC provides a practical computational strategy for improving the functional imaging performance of existing multiphoton microscopes without requiring fundamentally new optical hardware. In three-photon imaging, it enables a more temporally informative readout of deep-brain calcium dynamics while preserving spatial detail. In mesoscale two-photon imaging, it helps alleviate the speed penalty imposed by dense cellular-resolution sampling over large fields of view. In voltage imaging, it enables a more temporally informative 400-Hz readout on a standard resonant-galvo two-photon microscope and improves the recoverability of fast membrane-potential-related dynamics under the current experimental and analytical conditions. More broadly, our results suggest that reconstruction strategies tailored to specimen-matched structure and acquisition physics may substantially expand the functional operating range of scan-limited multiphoton microscopes. This framework may help enable tighter integration of acquisition design and computational recovery in future multiphoton imaging systems.

## Methods

### Multiphoton imaging system

The deep *in vivo* imaging was performed using a custom-built three-photon (3P) microscope. The excitation source was a 1,300 nm, 1 MHz repetition rate laser (I-OPA-TW-F, Light Conversion), with pulse dispersion compensated by a two-prism (SF11) compressor. A 25× water-immersion objective (XLPLN25XWMP2, NA 1.05, Olympus) was employed. Emitted fluorescence was epi-collected, separated by a 775 nm dichroic beam splitter (FF775-Di01-25×36, Semrock), filtered through a 520/44 nm band-pass filter, and detected using a GaAsP photomultiplier tube (PMT, H7422P-40, Hamamatsu). Scanning was achieved with a pair of galvanometer mirrors (6215H, Cambridge Technology). The PMT current was converted to voltage by a transimpedance amplifier (C12419, Hamamatsu).

The mesoscale two-photon (2P) calcium imaging and voltage imaging were conducted on a custom-built two-photon microscope. The excitation source was a tunable femtosecond laser (Chameleon Discovery NX, Coherent) set to 920 nm. For large-area scanning, a 4× objective (custom, NA 0.296) was used for a 6 × 6 mm² field of view, and a 16× objective (Nikon, NA 0.8) was used for a 1.5 × 1.5 mm² field of view. For voltage imaging, a 16× objective (Nikon, NA 0.8) was used, yielding a field of view of 167 × 714 μm^2^. Scanning was performed using a galvanometer (Y-axis, 6215H, Cambridge Technology) and a resonant mirror (X-axis, CRS8K, Cambridge Technology) via a telecentric telescope (LSM54-850, Thorlabs). Fluorescence detection utilized the same filter set, PMTs, and amplifiers as the 3P microscope.

For both microscopes, the sample was mounted on a motorized stage (MP-285A, Sutter Instrument). Image acquisition and stage control were managed by vDAQ and ScanImage software (Vidrio Technologies) running in MATLAB (MathWorks). All functional imaging was performed in bidirectional scanning mode without frame averaging. Key imaging parameters for each experiment are summarized in Supplementary Table 1. Anisotropic sparse sampling was achieved by reducing the number of lines scanned in the slow axis (e.g., from 512 to 128), while maintaining the fast-axis resolution and field of view, thereby proportionally increasing the imaging frame rate.

### Architectures and objective functions of SPARC

SPARC employs a unified two-stage neural network architecture that decomposes the upsampling reconstruction task in scan-limited multiphoton imaging into two consecutive subtasks: denoising and upsampling. The first stage retains maximum effective information to facilitate subsequent upsampling reconstruction. Specifically, the anisotropic sparse input *S* (*H*/4 × *W* ×*T*) is first split into two interleaved subsets based on odd and even frames (denoted as *S*_o_ and *S*_e_, each with a size of *H*/4 × *W* × *T*/2). The image stack of odd frames is first denoised by a 3D lightweight U-net^48^ (Supplementary Fig. 2), denoted as *F_denoised_* to stabilize and retain recoverable measurement information. The denoised odd-frame image stack and raw even frames are used to construct the denoising loss function (*L_denoised_*). We employ a linear combination of L1-norm loss and L2-norm loss as the loss function to optimize the parameters of the denoising module:

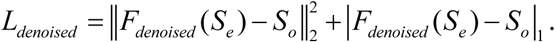

Subsequently, the output of Stage Ⅰ is fed into Stage Ⅱ (*F_upsampled_*) to recover the slow-axis information lost during the sampling process. The backbone of the deep-upsampling module mirrors that of the 3D U-net in the denoising module, but an upsampling convolution module is added after the decoder. Meanwhile, under the guidance of the high-resolution full-frame structural reference, *S*_e_ recovers the slow-axis information to a certain extent in the form of physics-aware weighted upsampling, denoted as *f_ref_*. This reference forms the deep-upsampling loss function (*L_upsampled_*) in conjunction with the output of Stage Ⅱ:

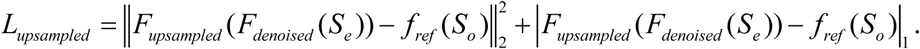

The parameters of the two-stage neural network are optimized through the joint backpropagation of the denoising loss and deep-upsampling loss:

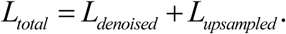

### Physics-aware weighted upsampling

In the deep-upsampling loss function of SPARC, we introduce a physics-aware weighted upsampling strategy to generate the upsampled anisotropic even-frame slices. Prior to fast multiphoton functional imaging, a set of full-frame reference stacks is acquired at a low temporal but high spatial resolution; the time-averaged image from this set serves as the structural reference. This reference, which covers the same field of view as the anisotropic even slices, provides a sample-specific spatial scaffold that constrains the recovery process by the intrinsic tissue structure. Specifically, column pixel patches in anisotropic even slices are upsampled by a factor of 4 based on the weight ratio derived from the corresponding blocks in the structural reference image. The values of adjacent pixels are then fused according to their spatial proximity weighting to generate 4× upsampled column blocks. Iterating this operation over the full extent of each anisotropic even slice yields the final upsampled targets used to compute the deep-upsampling loss term.

### Model training and inference

The SPARC model was trained on a workstation equipped with an Intel Xeon Gold 6248R CPU (3.00 GHz) and an NVIDIA A5000 GPU, with implementation based on Python 3.9.0 and PyTorch 1.8.0. Prior to model training, input data augmentation was performed to generate 50,000 3D image patches via random cropping, horizontal flipping, and vertical flipping. Specifically, the patches had spatial–temporal dimensions of 32 × 128 × 32 (height × width × frames) for calcium imaging and 30 × 128 × 32 for voltage imaging, which effectively expanded the size of the training dataset. The Adam optimizer was adopted, with an initial learning rate of 1 × 10^-4^, as well as exponential decay parameters of 0.9 for the first moment and 0.999 for the second moment. Model training was conducted with a batch size of 1 over 30 epochs, and convergence was generally achieved within approximately 8 hours. Upon convergence, the optimized model weights of the last epoch were loaded, allowing SPARC to perform noise suppression and anisotropic sampling reconstruction for scan-limited multiphoton imaging.

Similar to other algorithms, SPARC exhibits sample dependence: under the same imaging system, only one-time model training is required for imaging samples of the same type, and analogous samples can be directly reused in the trained model. During the training and inference phases, SPARC utilizes two fully independent datasets - acquired from the same mouse and the same imaging system but with different fields of view - for model training, generalization performance verification, and reconstruction effect evaluation. This strategy effectively eliminates performance assessment biases induced by overlapping data utilization.

Both the Self-vision and DFCAN algorithms were trained and performed isotropically downsampled input reconstruction using their default network architectures and hyperparameters. When processing anisotropically downsampled data, the Self-vision algorithm modifies the final upsampling layer of the network by adjusting the upsampling scaling factors correspondingly along the two dimensions. The datasets used for training and inference are completely identical to those of SPARC.

### Data simulation

Synthetic calcium imaging data were used for quantitative assessment of our SPARC method and comparative analysis against Self-vision and DFCAN. The simulation pipeline comprised synthesizing noise-free calcium imaging sequences (ground truth) and adding different levels of mixed Gaussian-Poisson noise. To generate noise-free calcium imaging data, we employed Neural Anatomy and Optical Microscopy (NAOMi)^35^ - a simulation method which was utilized to create realistic calcium imaging datasets for assessing multiphoton microscopy methods. The parameters of our simulation were listed in Supplementary Table 2; those not specified were set to default values. Simulated data exhibited highly consistent spatiotemporal features with experimentally acquired data, including key neuronal anatomy (such as cell bodies, neuropils, dendrites), neural activity, and blood vessels. For noise simulation, Poisson sampling was first performed on noise-free images to introduce content-dependent Poisson noise, followed by the addition of content-independent Gaussian noise to these preprocessed images. Poisson noise was designated as the dominant noise source.

### Data analysis of neuronal activity in calcium imaging

All acquired imaging data were preprocessed using a custom-written preprocessing pipeline. First, PMT ripple noise was automatically identified and excluded based on pixel intensity thresholds of < 700 a.u. (for 3P data) or 100 a.u. (for 2P data). Subsequently, a rigid motion correction algorithm was employed to mitigate in-plane motion artifacts across the entire imaging session. Regions of Interest (ROIs) corresponding to individual neurons were manually delineated and segmented via ImageJ (Fiji). The raw fluorescence time trace (*F*) for each ROI was extracted. The baseline fluorescence (*F*_0_) was defined as the average signal within the 10th to 50th percentiles of the raw fluorescence time trace’s intensity distribution. The relative fluorescence change was then calculated as Δ*F*/*F*_0_ = (*F* − *F*_0_) / *F*_0_.

Calcium events were detected based on the Δ*F*/*F*_0_ trace. A transient was identified as a calcium event if its peak amplitude exceeded a threshold of the mean + 1 × standard deviations of the entire Δ*F*/*F*_0_ trace, following established criteria. For each detected calcium event, the following kinetic parameters were quantified:

Amplitude (*A*_peak_): The peak Δ*F*/*F*_0_ value of the event.

For kinetic comparisons between the 13.52-Hz SPARC signal and the 3.38-Hz-equivalent signal, calcium transients were analyzed using a matched-event strategy. Events were first identified in the higher-temporal-resolution trace, and the corresponding time points were mapped to the temporally matched low-frame-rate trace to compare the apparent kinetics of the same underlying transients.

Rise Time: The duration from event onset to peak. For each matched event, the onset time (*t*_start_) was defined as the last time point before the peak (*t*_peak_) at which the signal fell below baseline (*F*_0_), and rise time was calculated as *t*_rise_ = (*t*_peak_ − *t*_start_) × Δ*t*.

Decay Half-Time: The duration required for the signal to decay from its peak to 50% of the peak amplitude. For each matched event, the half-decay time point (*t*_half_) was defined as the first time point after the peak at which the signal fell below 0.5 × *A*_peak_, and decay half-time was calculated as *t*_1/2_ = (*t*_half_ − *t*_peak_) × Δ*t*.

Data points that met predefined exclusion criteria were removed from the analysis.

### Data analysis in voltage imaging

We manually selected ROIs from the averaged images of the registered image sequence. A rolling percentile filter (50%, 10 s window) was applied to the mean-intensity trace of the ROI to derive the fluorescence baseline *F*_0_ of voltage trace. In addition to Δ*F*/*F*_0_ traces, we calculated SNR traces as the ratio between Δ*F* (functional change) and √*F* (Poisson noise). For detected-spike analysis, the SNR traces were further subjected to a 100-Hz 12th order low-pass Butterworth filter. Detected spikes were defined as events crossing a threshold of SNR = mean + 2 × standard deviations under the current analysis pipeline.

Power spectral density (PSD) characterizes the distribution of signal power across frequency bands, illustrating energy allocation among distinct frequency components. This metric is widely used to assess intrinsic neuronal oscillations, stimulus-evoked dynamics, and background noise profiles. To suppress high-frequency random noise, averaged PSD curves of population voltage activity were computed via Welch’s periodogram averaging method in MATLAB. A Hamming window was implemented to minimize spectral leakage.

To quantify the synchronization between the phase of neuronal voltage activity and the timing of air-puff stimulation, we adopted the phase locking value (PLV). Specifically, neuronal Δ*F*/*F*_0_ signals were first bandpass-filtered (0.1–10 Hz) using a 4th-order Butterworth zero-phase filter to retain relevant frequency components. Subsequently, the analytic signal was obtained via the Hilbert transform, and the instantaneous phase was extracted as the argument of this analytic signal.

We then computed the PLV based on the neuronal phase at a 0.3-s delay relative to the stimulation onset. The PLV is defined as:

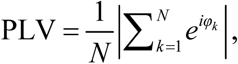

where *φ_k_* denotes the neuronal phase corresponding to the *k*-th stimulation trial, and *N* represents the total number of valid stimulation events.

### Performance metrics

To rigorously evaluate the reconstruction performance of the various algorithms, we employed a comprehensive set of quantitative metrics. Given a reconstructed image stack *S_x_* and its corresponding ground truth *S_y_*, the evaluation criteria are defined as follows.

The Structural similarity index (SSIM) quantifies perceptual similarity by evaluating three key components: luminance, contrast, and structural consistency. The definition is:

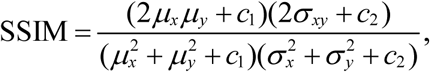

where {*μ_x_*, *μ_y_*} and {*σ_x_*, *σ_y_*} are the means and variances of *S_x_* and *S_y_*, respectively. *σ_xy_* is the covariance of *S_x_* and *S_y_*. The two constants *c*_1_ and *c*_2_ are defined as *c*_1_ = (*k*_1_*L*)^2^ and *c*_2_ = (*k*_2_*L*)^2^ with *k*_1_ = 0.01, *k*_2_ = 0.03 and *L* = 65,535.

The mean absolute error (MAE) serves as a spatial metric that measures the average absolute deviation between the reconstructed image stack and the ground truth, thereby providing a direct assessment of the overall reconstruction error. The MAE between *S_x_* and *S_y_* is defined as:

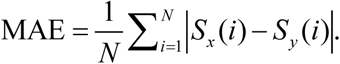

Pearson’s correlation coefficient (r) was utilized as a temporal metric to characterize the similarity between the reconstructed traces and the ground-truth traces. The Pearson correlation between signal *S_x_* and reference signal *S_y_* is defined as:

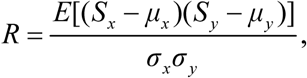

where *E* represents the arithmetic mean. {*μ_x_*, *μ_y_*} and {*σ_x_*, *σ_y_*} are the means and variances of *S_x_* and *S_y_*, respectively.

The SNR for each neuronal Δ*F*/*F*_0_ time series was computed using a robust estimation method based on the median absolute deviation (MAD). The signal (*S*) is estimated as the difference between the 95th percentile (*P*_95_) and the median of the Δ*F*/*F*_0_ trace, i.e.,

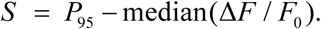

This measure reflects the typical amplitude of calcium transients while being robust to outliers.

The noise (*N*) is estimated from the median of absolute differences between consecutive time points. First, the median absolute deviation of the first-order difference was computed:

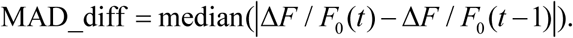

This value was then scaled by a factor of 1/0.6745 to obtain a consistent estimator of the standard deviation under a normal distribution:

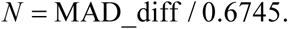

Finally, the SNR in decibels (dB) was calculated as:

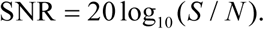

### Animals

All animal procedures were performed in accordance with protocols approved by the Institutional Animal Care and Use Committee of the Department of Laboratory Animal Science at

Fudan University (Protocol #2022JS-ITBR-013). Mice were housed under a 12-hour light/dark cycle with ad libitum access to food and water.

The following mice were used for different imaging experiments. Three-photon deep imaging was performed in an adult C57BL/6 mouse expressing GCaMP6f in the sensory cortex via viral injection. Mesoscale two-photon structural imaging was performed in an adult CaMKIIα::Ai162 transgenic mouse expressing GCaMP6s predominantly in cortical excitatory neurons. Mesoscale two-photon functional imaging was performed in an adult C57BL/6 mouse expressing GCaMP6f in the cortex via viral injection. Two-photon voltage imaging was performed in an adult C57BL/6 mouse expressing the voltage sensor ASAP5-Kv in the cortex via viral injection.

### Viral injection and cranial window implantation

Mice were anesthetized with 1–2% isoflurane and fixed in a stereotactic frame on a heating pad. Ophthalmic ointment was applied to prevent corneal drying. Viral injections were performed using a glass micropipette connected to a 10 μL Hamilton syringe mounted on a microinjection pump. The virus was injected at a rate of 50 nL / min, and the pipette was left in place for 10 minutes post-injection before withdrawal to minimize backflow.

For three-photon imaging, 200 nL of AAV2/9-CaMKIIα-GCaMP6f (∼1.55×10^13^ vg/mL, Obio) was injected into the sensory cortex (AP: −1.5 mm, ML: +1.5 mm, DV: −0.6 mm). For two-photon mesoscale functional imaging, 200 nL of the same AAV2/9-CaMKIIα-GCaMP6f virus was injected into the cortex (AP: −1.06 mm, ML: +1.5 or +2.0 mm, DV: −0.3 mm). For two-photon voltage imaging, 200 nL of AAV-CaMKIIα-ASAP5-Kv (∼2.8×10¹³ vg/mL, Obio) was injected into the cortex (AP: −1.0 mm, ML: +0.9 mm, DV: −0.3 mm).

Following a 3-week post-injection recovery and expression period, a chronic cranial window was implanted. A circular craniotomy (4 mm diameter for most experiments, 7 mm for the largest mesoscale imaging) was made using a dental drill, leaving the dura intact. The bone flap was carefully removed, and any bleeding was controlled with an absorbable hemostatic agent. A custom-sized glass coverslip (4 mm or 7 mm diameter, 150 μm thick) was placed over the brain and sealed with tissue adhesive. Finally, a custom titanium headplate was attached to the skull using cyanoacrylate adhesive to facilitate head-fixation during imaging.

### Behavioral paradigm

Prior to imaging, mice were acclimated to head fixation under the microscope. Each mouse was head-fixed and allowed to habituate to the experimental setup for 30 minutes per day over 5 consecutive days.

For the voltage imaging experiments involving air-puff stimulation, a controlled air-puff delivery system was used. Compressed air was gated by a three-way solenoid valve, which was triggered by a TTL signal from a data acquisition device. The TTL signals were generated by a custom MATLAB script synchronized with the microscope frame clock from vDAQ, ensuring precise temporal coordination between stimulus delivery and image acquisition. The air puff was directed toward the mouse’s contralateral whisker pad.

### Statistical analysis and quantification

All data processing and statistical analyses were performed using custom-written scripts in MATLAB. Data visualization formats and detailed statistical parameters including the sample size, type, and replicate number are explicitly specified in the corresponding figure legends.

## Resource Availability

### Lead contact

Further information and requests for resources and reagents should be directed to and will be fulfilled by the lead contact, Bo Li.

### Materials availability

This study did not generate new unique reagents.

### Data and code availability

The data that support the findings of this study are available from the corresponding author upon reasonable request. The customized code of SPARC is available at https://github.com/ShoupeiLiu/SPARC-master. The other algorithms mentioned as follows: Self-vision software is installed from https://github.com/frankheyz/s-vision; DFCAN software is installed from https://github.com/qc17-THU/DL-SR.

## Acknowledgements

This work was supported in part by the Brain Science and Brain-like Intelligence Technology – National Science and Technology Major Project (2025ZD0216500), the National Natural Science Foundation of China (32471142), the Science and Technology Commission of Shanghai Municipality (20JC1419500), and the Shanghai Pilot Program for Basic Research – Fudan University (21TQ1400100, 22TQ019).

## Author Contributions

B.L. and Y.H.Y. conceived and supervised the project. S.L. developed the reconstruction algorithm. J.H.H. and S.L. generated the simulated datasets and benchmarked the existing reconstruction methods. Y.G.Z. and Y.F.Z. performed the animal surgeries. Y.G.Z. and S.L. performed the imaging experiments and analyzed the data. B.L., S.L., Y.G.Z., and Y.H.Y. wrote the manuscript. B.L. provided funding support.

