## Supplemental Figure 1-3, Table 1-2 for "Structure-guided, physics-aware reconstruction expands scan-limited multiphoton imaging"

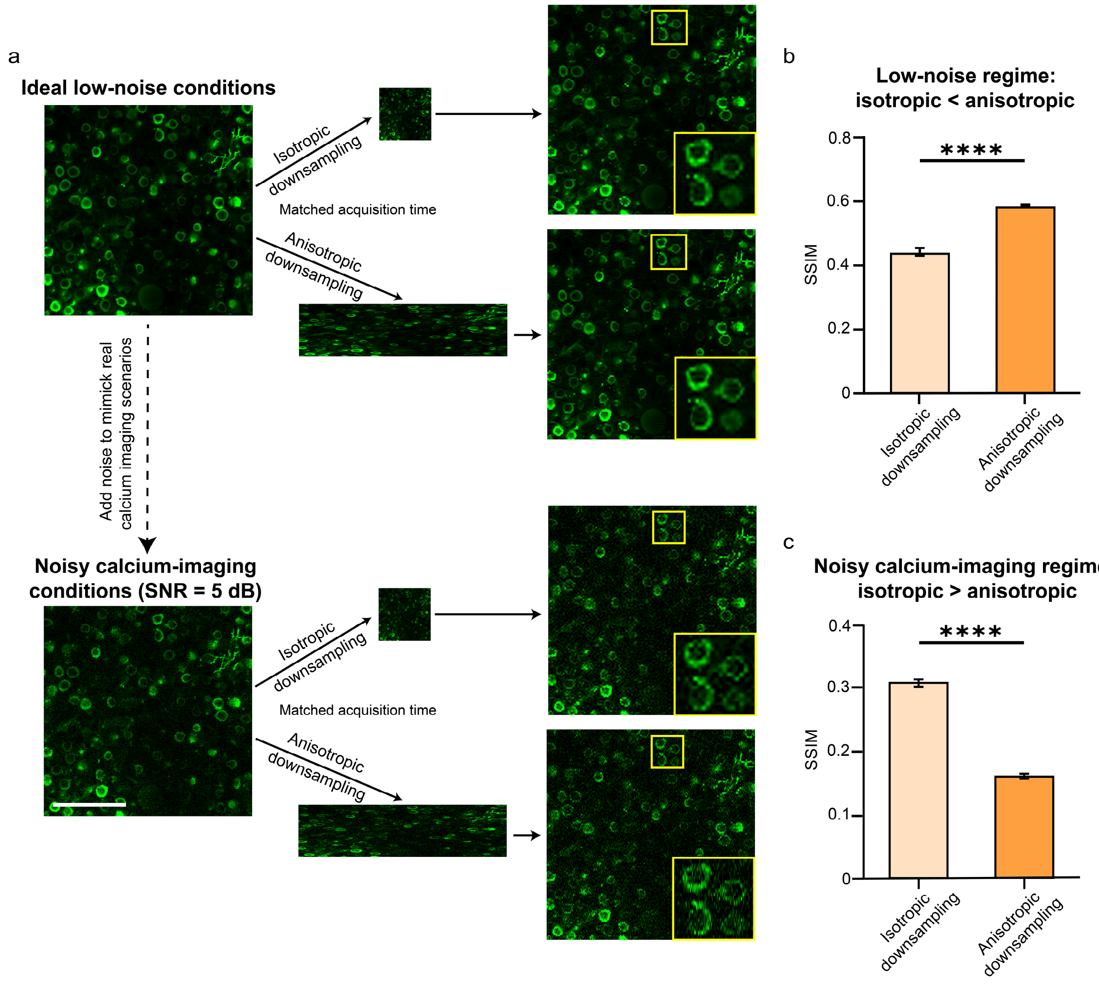


**Supplementary Fig. 1 Noise-dependent advantage of anisotropic versus isotropic sampling under matched acquisition time. a**, Flowchart and representative single-frame reconstructions for isotropic and anisotropic downsampling using Self-vision under low-noise and noisy calcium-imaging conditions. Yellow boxes indicate magnified regions. Scale bar, 100 μm. **b**, Structural similarity index (SSIM) of reconstruction from isotropic and anisotropic downsampling under low-noise conditions, showing superior performance for anisotropic inputs. **c**, SSIM of reconstruction from isotropic and anisotropic downsampling under noisy calcium-imaging conditions, showing superior performance for isotropic inputs. Error bars indicate s.d. Statistical significance was assessed using a paired $t$-test ($n$ = 1008 frames). ****$P < 0.0001$. Self-vision was used here as a representative example to illustrate how the relative performance of isotropic and anisotropic sampling depends on noise conditions.

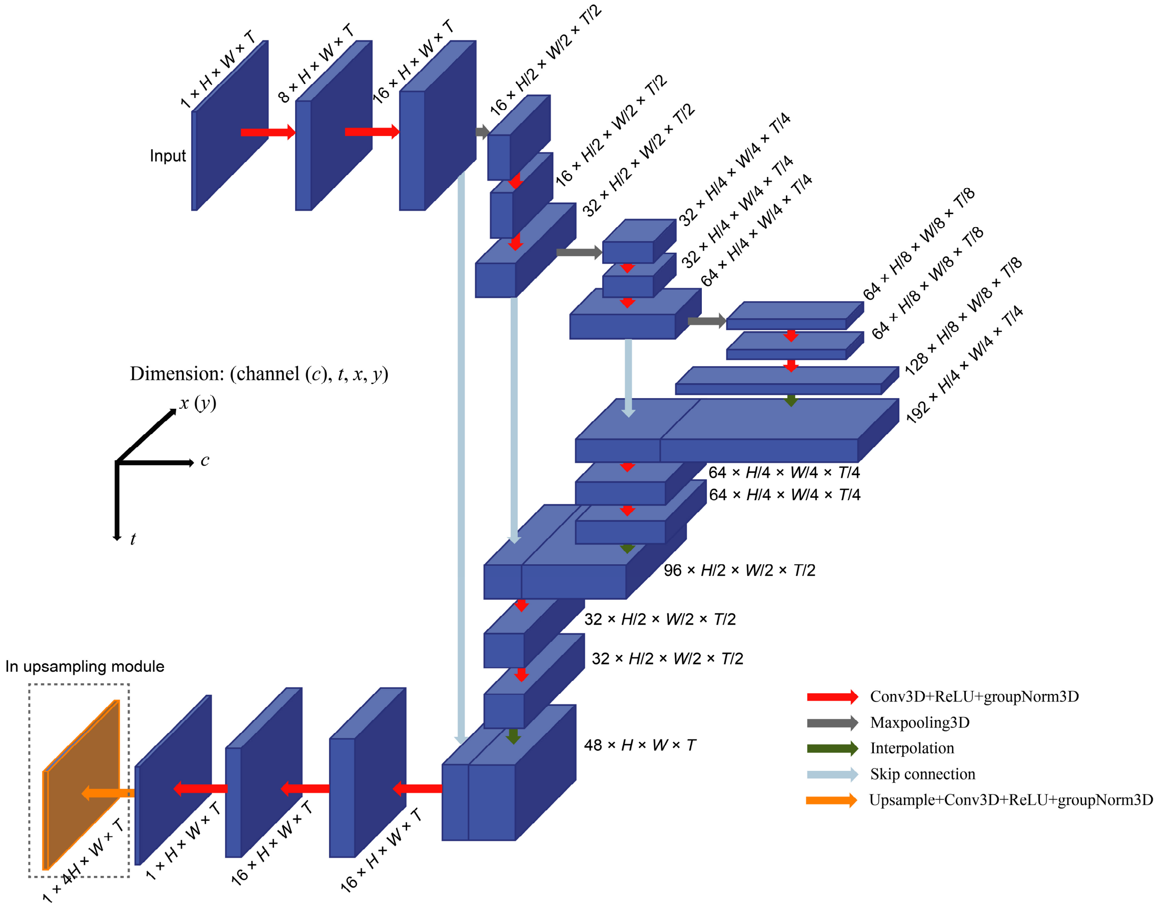


**Supplementary Fig. 2 Network architecture.** A simplified 3D U-Net was used as the backbone architecture of SPARC, comprising a 3D encoder, a 3D decoder, and skip connections between them. In the upsampling stage, an additional upsampling module is appended after the 3D U-Net to recover information lost along the slow axis.

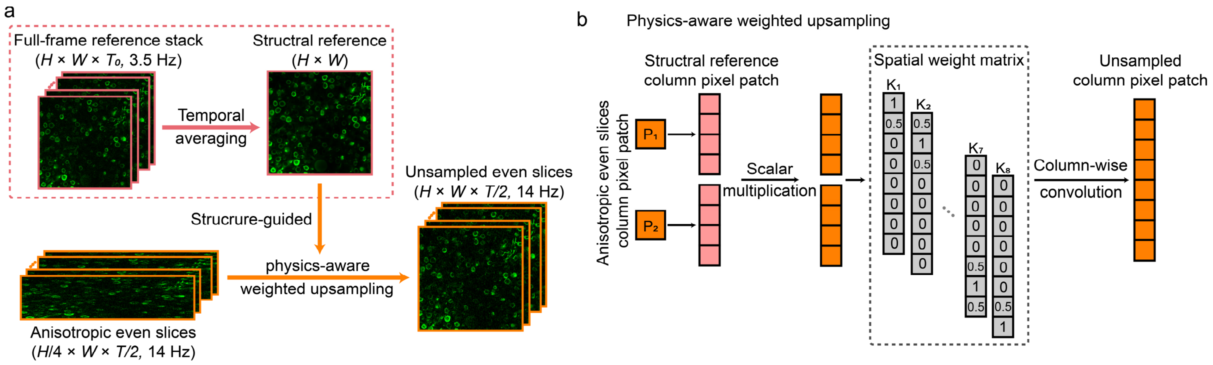


**Supplementary Fig. 3 Conceptual diagram of sample-specific structural reference and physics-aware weighted upsampling. a**, Generation of upsampled anisotropic even slices. During SPARC training, a sample-specific structural reference is obtained by temporally averaging a low-frame-rate full-frame reference stack. This structural reference is then used to guide physics-aware weighted upsampling of the original anisotropic even slices, generating high-resolution target slices for loss computation. **b**, Conceptual diagram of physics-aware weighted upsampling. Single column pixels in a sparse anisotropic slice are first upsampled by a factor of 4 based on the weight ratio derived from the corresponding blocks in the structural reference image. Adjacent pixels are then fused according to their spatial distance distribution to generate 4× upsampled column blocks. Applying this procedure to the full anisotropic slice yields the reference-guided training targets used for deep-upsampling loss computation

### Supplementary Table 1. List of imaging parameters

| Imaging data | Fluorescence | Image size (pixels) | FOV ($\mu m^2$) | Frame rate (Hz) | Objective | Sampling mode |
|---|---|---|---|---|---|---|
| Fig. 3a | GCaMP6f | 512 × 512 | 285 × 285 | 3.38 | Olympus 25x | Conventional full frame |
| | GCaMP6f | 128 × 512 | 285 × 285 | 13.52 | Olympus 25x | Anisotropic sparse |
| Fig. 4a | GCaMP6s | 4096 × 4096 | 6000 × 6000 | 3.8 | Meso2P 4×, | Conventional full frame |
| | GCaMP6s | 1024 × 4096 | 6000 × 6000 | 15.21 | Meso2P 4×, | Anisotropic sparse |
| Fig. 4d | GCaMP6f | 1024 × 4096 | 1500 × 1500 | 15.21 | Nikon 16x | Anisotropic sparse |
| Fig. 5b | ASAP5 | 120 × 512 | 167 × 714 | 100 | Nikon 16x | Conventional |
| | ASAP5 | 30 × 512 | 167 × 714 | 400 | Nikon 16x | Anisotropic Sparse |

Notes:

Image size refers to the pixels per frame, which is in unit of pixels, height × width (slow axis × fast axis).

FOV refers to the actually-illuminated and sampled scanning region.

Meso2P 4× is an in-home developed air objective with a NA of 0.3, designed specifically for mesoscopic two-photon imaging.

**Supplementary Table 2. Key imaging parameters for NAOMi-generated images**

| **Physiological parameters** | | **Imaging parameters** | |
|---|---|---|---|
| Sample volume | 500 ×500 × 50 $\mu m^3$ | FOV | 250 × 250 $\mu m^2$ |
| Average neuron radius | 5.9 μm | Pixel size | 0.488 μm |
| Number of neurons | 1250 | Image size | 512 × 512 pixels |
| Average firing rate | 0.25 | Frame rate | 14 Hz |
| Vasculature simulation | ON | Imaging depth | 300 μm |
| Background dendrites | ON | Excitation NA | 0.6 |
| Calcium indicator | GCaMP6 | Detection NA | 0.8 |
| Fluorophore concentration | 10 μM | Excitation power | 50 mW |
| **Laser source** | | Number of frames | 1008 |
| Wavelength | 920 nm | Objective focal length | 4.5 mm |
| Repetition frequency | 80 MHz | Brain motion | OFF |
| Pulse width | 150 fs | PSF type | Gaussian |
